# Single-nucleus RNA-seq resolves spatiotemporal developmental trajectories in the tomato shoot apex

**DOI:** 10.1101/2020.09.20.305029

**Authors:** Caihuan Tian, Qingwei Du, Mengxue Xu, Fei Du, Yuling Jiao

## Abstract

Single cell transcriptomics is revolutionizing our understanding of development and response to environmental cues^1–3^. Recent advances in single cell RNA sequencing (scRNA-seq) technology have enabled profiling gene expression pattern of heterogenous tissues and organs at single cellular level and have been widely applied in human and animal research^4,5^. Nevertheless, the existence of cell walls significantly encumbered its application in plant research. Protoplasts have been applied for scRNA-seq analysis, but mostly restricted to tissues amenable for wall digestion, such as root tips^6–10^. However, many cell types are resistant to protoplasting, and protoplasting may yield ectopic gene expression and bias proportions of cell types. Here we demonstrate a method with minimal artifacts for high-throughput single-nucleus RNA sequencing (snRNA-Seq) that we use to profile tomato shoot apex cells. The obtained high-resolution expression atlas identifies numerous distinct cell types covering major shoot tissues and developmental stages, delineates developmental trajectories of mesophyll cells, vasculature cells, epidermal cells, and trichome cells. In addition, we identify key developmental regulators and reveal their hierarchy. Collectively, this study demonstrates the power of snRNA-seq to plant research and provides an unprecedented spatiotemporal gene expression atlas of heterogeneous shoot cells.

---

Plant aerial tissues and organs are generated from the shoot apical meristem (SAM), which embraces a central zone (CZ) of stem cells, an organizing center (OC) beneath the CZ, and a peripheral zone (PZ) surrounding the CZ^11^. The OC serves as the stem cell niche by providing stem cell-promoting cues. Some of the stem cell progenies are displaced from the CZ into the PZ, where they form organ primordia. The zonation into functional domains in the SAM is dynamic, as shown by molecular markers.

The SAM and early leaf primordium have a complex cellular architecture consisting of heterogeneous cell types embedded in cell walls of different composition. Outside the epidermis layer, cuticle and wax covering helps to reduce water loss but also impedes enzymatic dissociation of cells. To dissociate protoplasts from the shoot apex, prolonged intensive enzymatic digestion is required. Wide-spread ectopic activation^12^ and stochastic gene expression^13^ are associated with protoplasting, in which cell walls are digested. To estimate the effects of protoplasting on gene expression, we used fluorescent reporters to monitor the gene expression in *Arabidopsis thaliana* leaves and mesophyll protoplasts, and observed frequent ectopic activation. For example, *WUSCHEL-RELATED HOMEOBOX 2* (*WOX2*) is not expressed in leaves (Extended Data Fig. 1a). We followed over 10,000 protoplasts and observed that over 21% of the protoplasts expressed *WOX2* (Extended Data Fig. 1b). This observation strongly suggests the existence of ectopic random activation of gene expression by protoplasting.

To circumstance of the cell wall barriers, ectopic gene expression artifacts, and depletion of certain cell types following protoplasting, we established a plant tissue processing pipeline to isolate high-quality nuclei, which are compatible with high-throughput snRNA seq. We applied the tissue processing and snRNA-seq pipeline to interrogate cell-type diversity and spatiotemporal developmental trajectories of vegetative tomato shoot apex tissues, which have thicker cell walls than *Arabidopsis* and are more resistant to protoplasting. We dissected tomato shoot apices under dissection microscope and retained the SAM region and early leaf primordia up to P_3_, which denotes the third youngest leaf primordium, from 2-week-old plants (Extended Data Fig. 2). We established a pipeline to efficiently remove cell wall debris and plastids, while retaining normal nuclei morphology (Extended Data Fig. 3, and see Methods). Purified nuclei with high quality were sent for encapsulation by the droplet-based 10x Genomics platform (Fig. 1a). After quality control and filtration, we obtained 13,377 nuclei and detected the expression of 21,402 genes, which corresponds to 62.8% of annotated genes. To evaluate the reproducibility and sensitivity of the snRNA-seq data, we performed bulk RNA-seq with comparable shoot apices. Pooled snRNA-seq detected (FPKM > 1) 92.3% of genes detected by bulk RNA-seq, suggesting high sensitivity of snRNA-seq. Furthermore, the gene expression profiles of pooled snRNA-seq and bulk RNA-seq are highly correlated (*r* = 0.90, Spearman correlation coefficient; Fig. 1b), indicating high reproducibility.

**Fig. 1:**
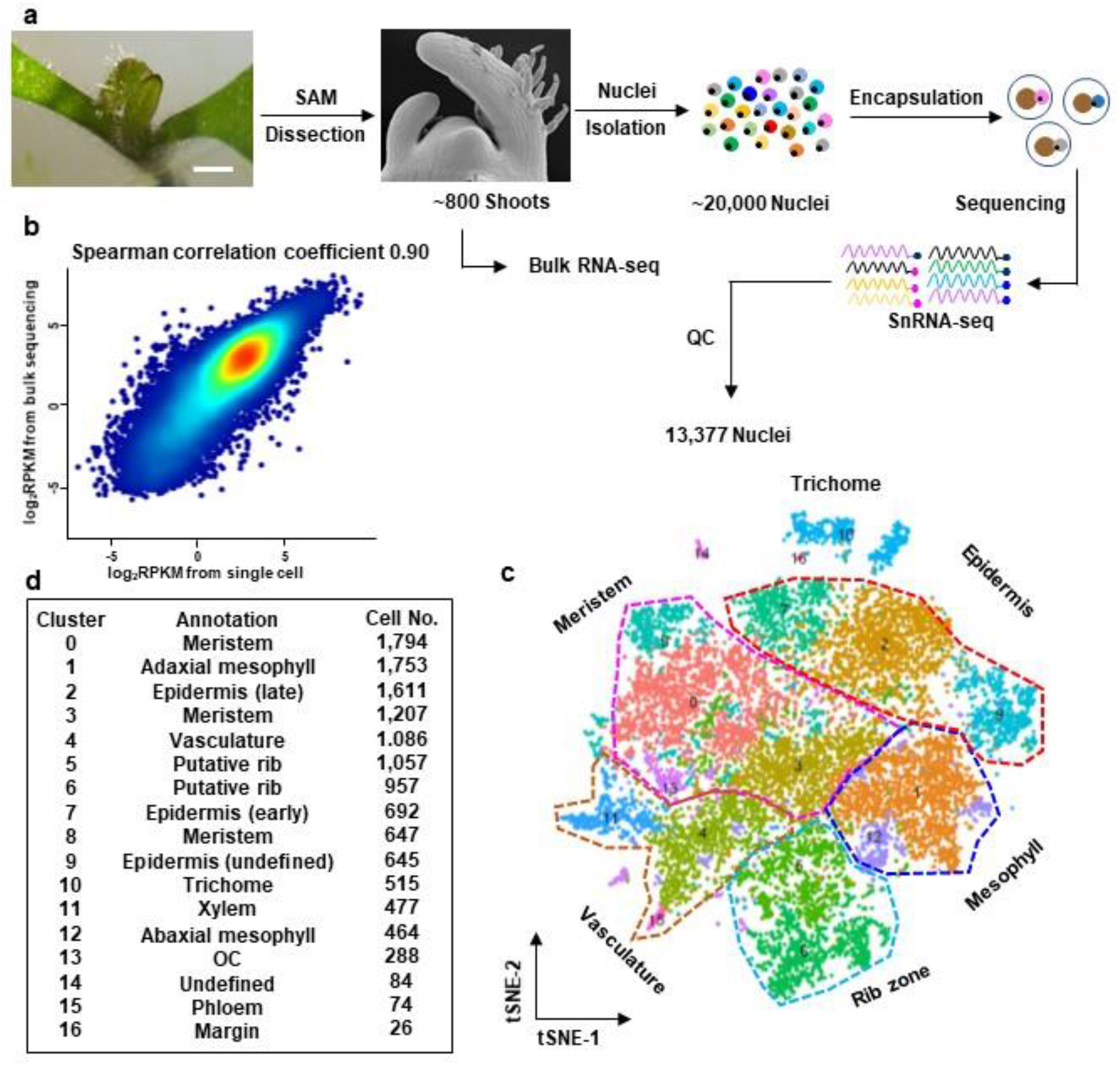
A cell atlas of tomato shoot apex by snRNA-seq. **a,** Procedure of snRNA-seq. **b**, Correlation between snRNA-seq and bulk RNA-seq. **c**, Cell clusters displayed by *t*-SNE. **d**, Cell cluster annotation and identified number of cells in each cluster.

To identify distinct cell type populations, we applied unsupervised clustering analysis. The scaled data was reduced into 20 approximate principal components (PCs) by linear dimensional reduction. The *t*-distributed stochastic neighborhood embedding (*t*-SNE) algorithm was employed and grouped the nuclei into 16 cell clusters (Fig. 1c). Each cluster possessed a remarkable differential gene expression pattern (Extended Data Fig. 4). A series of specific marker genes for each cluster were identified (Fig. 2a, Extended Data Fig.5, Extended Data Table 1). To annotate these clusters, we correlated the cluster marker genes with known markers. In addition, we correlated with genes whose *Arabidopsis* homologous genes are enriched in corresponding cell domains^14–17^. We found significant enrichment of homologous genes to *Arabidopsis* cell domain-specific genes, including epidermis, mesophyll, and vasculature domains, in several clusters (Extended Data Fig. 6). In addition, putative orthologs of the *Arabidopsis* meristem marker genes *Sl SHOOT MERISTEMLESS* (*SlSTM*), and *Sl BREVIPEDICELLUS* (*SlBP*), which are broadly expressed in the SAM, were specifically expressed in clusters 0, 3, 4, 5, 6, 7, 8 and 13 (Extended Data Fig. 5). This observation is consistent with the fact that a significant proportion of our samples correspond to the SAM. Based on the combination of the above information, we annotated the cell clusters into four cluster clouds, corresponding to epidermis and trichomes (clusters 2, 7, 9, 10 and 16), mesophylls (clusters 1, 12 and 14), vasculature (clusters 4, 11 and 15), and meristem cells (clusters 0, 3, 5, 6, 8 and 13) (Fig. 1c,d).

**Fig. 2:**
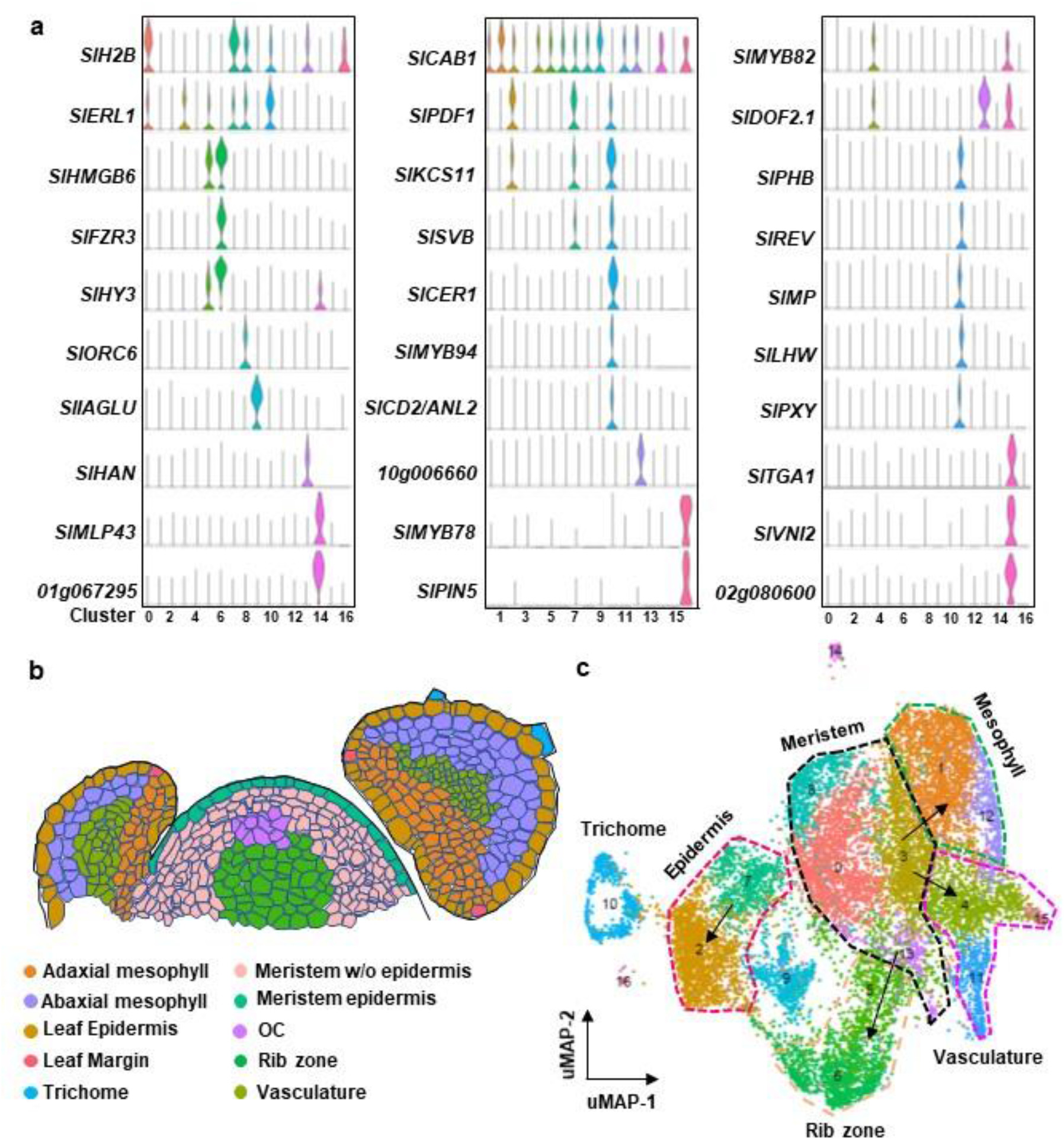
Cell heterogeneity and spatial distribution. **a**, Marker gene expression pattern in each cluster. Gene IDs are provided in Extended Data Table 2. **b**, Spatial distribution of different clusters in the shoot apex. **c**, Visualization of tomato shoot apex cell clusters by uMAP algorithm. Arrows indicate putative differentiation routes.

Clusters 1 and 12 comprise leaf primordium adaxial and abaxial cells, respectively (Fig. 1c, Extended Data Fig. 6). Consistent with future photosynthesis activity, genes involved in photosynthesis, such as *Sl CHLOROPHYLL A/B BINDING PROTEIN 1* (*SlCAB1*), and *Sl PHOTOSYSTEM I LIGHT HARVESTING COMPLEX GENE 2* (*SlLHCA2*), are remarkably enriched in cluster 1. Cluster 12 is enriched with genes responding to abiotic stimuli, as shown by gene ontology (GO) analysis (Extended Data Table 3), in addition to abaxial identity genes.

Cluster 4 comprises vasculature cells. In this cluster, genes homologous to *Arabidopsis* vasculature cell-specific genes are enriched, as well as *Sl DOF PROTEIN 2.1* (*SlDOF2.1*), a putative vasculature marker^18^. A neighboring cluster (11) comprises developing xylem cells with orthologs to *Arabidopsis* xylem identity markers, including *Sl PHLOEM INTERCALATED WITH XYLEM* (*SlPXY*), *Sl PHABULOSA* (*SlPHB*), *Sl CORONA* (*SlCNA*), *Sl LONESOME HIGHWAY* (*SlLHW*), *Sl MONOPTEROS* (*SlMP*), and *Sl REVOLUTA* (*SlREV*)^19^, are enriched. Another small neighboring cluster (15) corresponds to developing phloem cells and expresses orthologs of *Arabidopsis* phloem markers *Sl PHLOEM PROTEIN 2 A1 and B12* (*SlPP2-A1, SlPP2-B12*)^19^.

Clusters 2, 7 and 10 comprise epidermal cells with the expression of orthologs to *Arabidopsis* marker genes *Sl MERISTEM LAYER 1* (*SlML1*), *Sl PROTODERMAL FACTOR 2* (*SlPDF2*), *SlPDF1*^20^. Orthologs to cuticle development genes, such as *Sl FIDDLEHEAD* (*SlFDH*), *Sl FACELESS POLLEN 1* (*SlFLP1*), and *Sl PERMEABLE LEAVES3* (*SlPEL3*), are also expressed in these clusters. Cluster 7 contains fast dividing epidermal cells and is enriched with cell cycle S-phase genes, including histone genes and genes involved in chromatin replication. Putative meristem epidermis cells, which express *SlCLVATA3*, are mixed with other fast dividing epidermis cells, which we collectively termed as Epidermis (early). By contrast, cells in cluster 7 lack cell cycle gene expression and were termed Epidermis (late). Cluster 10 contains maturing trichome cells and expresses orthologs to *Arabidopsis* trichome specification genes, including *Sl ANTHOCYANINLESS 2* (*SlANL2*), and *Sl MIXTA* (*SlMX1*). Cluster 9 has overlapping expression patterns with *Arabidopsis* epidermal cells^14,16^, but remains different from above-mentioned clusters, representing a novel epidermal subgroup. Cluster 16 corresponds to leaf primordium margin cells, which are fast dividing with unique margin genes.

Clusters 0, 3, 5, 6, 8 and 13 comprise SAM cells and are enriched with cell cycle genes. The expression pattern of cluster 13 shows similarity to *Arabidopsis* OC and subepidermis (L2) cells of the inflorescence meristem, suggesting OC identity. Clusters 5 and 6 are enriched with G2-phase cell cycle genes, whereas clusters 0 and 8 are enriched with S-phase cell cycle genes, including DNA replication and chromatin modulation related genes. Together, we identified major cell types of the shoot apex, which provided expression information to the spatial distribution of shoot apex cells (Fig. 2b).

Single cell transcriptomics can capture cells with transition state, enabling us to trace the development trajectory of a specific cell type. To obtain a pandemic view of shoot apex cell developmental trajectories, we applied the uniform manifold approximation and projection (uMAP) algorithm to cluster and visualize the hierarchical structures of cell clusters (Fig. 2c). Whereas similar cell clusters were identified as compared with *t*-SNE, the clusters corresponding to meristem cells were located in the center. Continuous trajectories of shoot cell differentiation rooted to the meristem cells lead to epidermal cells, trichome cells, mesophyll cells, and vasculature cells.

To reconstruct developmental trajectories for key cell types, we used Monocle 2 to carry out pseudotime analysis. After leaf initiation at the PZ of the SAM, leaf mesophyll cells differentiate with a distinction between the adaxial and abaxial sides^21^, which is accompanied by vasculature formation^19^. We subjected clusters 1, 3, 4, 11, 12 and 15 to unsupervised pseudotime trajectory reconstruction and assembled developmental trajectories containing five branches (Fig. 3a,b). Cluster 3 meristem cells were assigned as the beginning of pseudotime. At the first branch, mesophyll cells (clusters 1 and 12) were clearly separated from vasculature cells (clusters 4, 11 and 15), which were subsequently separated into xylem cells and phloem cells at the next branch. Notably, another small group of vasculature cells separated from mesophyll cells, suggesting transdifferentiation of mesophyll progenitor cells into high-order vasculature. Cells in the first branching point of pseudotime express genes involved in auxin signaling, such as orthologs of *MP* and *PIN1* (Fig. 3c,d), which is consistent with the roles of auxin in leaf initiation^11^. Distinct gene expression patterns emerge along both differentiation trajectories. In the mesophyll branch, photosynthesis genes start to expression, as well as leaf abaxial and middle domain identity genes *SlFIL* and *SlKAN2*. In the vasculature branch, many vascular identity genes are expression. The commencement of *SlREV, SlLHW*and *SI DOF AFFECTING GERMINATION 1* (*SlDAG1*) orthologs is followed by *SlCNA, SlDOF5.6, SlPHB*, and *SlWOX4* orthologs, suggesting hierarchical regulation. The vasculature branch subsequently further branches into pro-phloem cells and pro-xylem cells. Cytokinin signaling is activated during pro-phloem specification, whereas auxin signaling and polar transport are activated during pro-xylem specification (Extended Data Fig. 7). Furthermore, we reconstructed a gene regulatory network (GRN) showing the complex regulation among transcription factors along the pseudotime (Extended Data Fig. 7d).

**Fig. 3:**
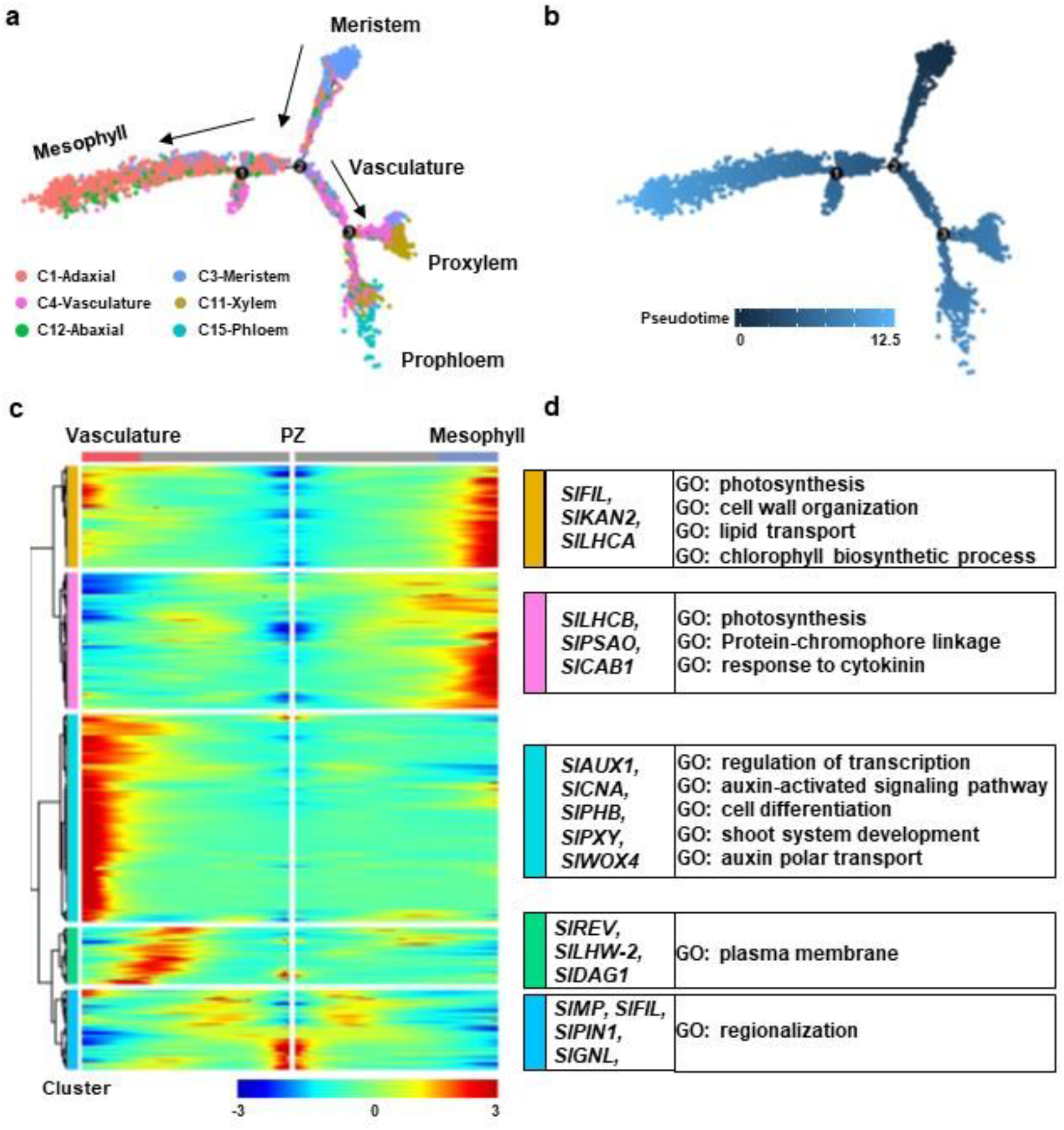
Developmental trajectory involved in leaf initiation and vasculature specification. **a** and **b,** Developmental trajectory of mesophyll and vasculature cells highlighting clusters (**a**) and pseudotime (**b**). **c**, Heatmap of branch-dependent gene expression patterns over the pseudotime. **d**, Marker genes and enriched GO terms of branch-dependent genes as shown in **c**.

The epidermis is a single layer of clonally related cells^20,22^, for which we analyzed in detail. Although the shoot apices contain only early trichome cells, snRNA-seq identified sufficient trichome cells for further analysis. Tomato displays multiple types of trichomes that can be divided into glandular and non-glandular types^23,24^. Although there is a lack of marker genes for glandular and non-glandular trichomes, we detected two subclusters in cluster 10 (Figure 1c, Extended Data Fig.8). Genes regulating glandular trichome formation, such as *SlMX1, SlWOOLLY* and *SlSVB* are enriched in putative glandular subcluster (subcluster 2), while genes regulating cuticle development, such as *SlCD2/ANL2, SlCSLA9, SlFDH* et al., were enriched in the other subcluster (subcluster 1) (Extended Data Fig. 8a,b). GO analysis showed that genes with higher expression in subcluster 1 are enriched with “Cell wall organization”, “Carboxylic acid biosynthetic process”, “Response to salt stress”, and “Response to ABA” terms, while in subcluster 2 are enriched with “Tissue Development” and “Lipid metabolic process”, indicating a role in early differentiation (Extended Data Table 4) Unsupervised pseudotime developmental trajectory analysis revealed that trichome cells are separated from other epidermis cells at the first branching point. Whereas branches toward mature epidermal cells are enriched with genes function in photosynthesis, cell wall organization, and organ morphogenesis, branches toward trichome cells are enriched with genes function in cell wall loosening, cuticle development, wax biosynthesis, cell morphogenesis, and response to stimulus (Extended Data Table 5).

We further reconstructed GRN using transcription factors with differential expression along pseudotime of the epidermis developmental trajectory. The GRN is centered at *SlPDF1* and *SlSVB*, and connects genes responsible for leaf initiation and leaf polarity, suggesting early interactions of these developmental programs (Fig. 4d, Extended Data Fig. 8b). Furthermore, both positive and negative regulators of trichome specification, such as *SlMX1, SlANL2, SlSPL8* and *SlNOK*, reside within interconnected distal ends of the GRN. Multiple feedback and feedforward loops are identified in the GRN.

**Fig. 4:**
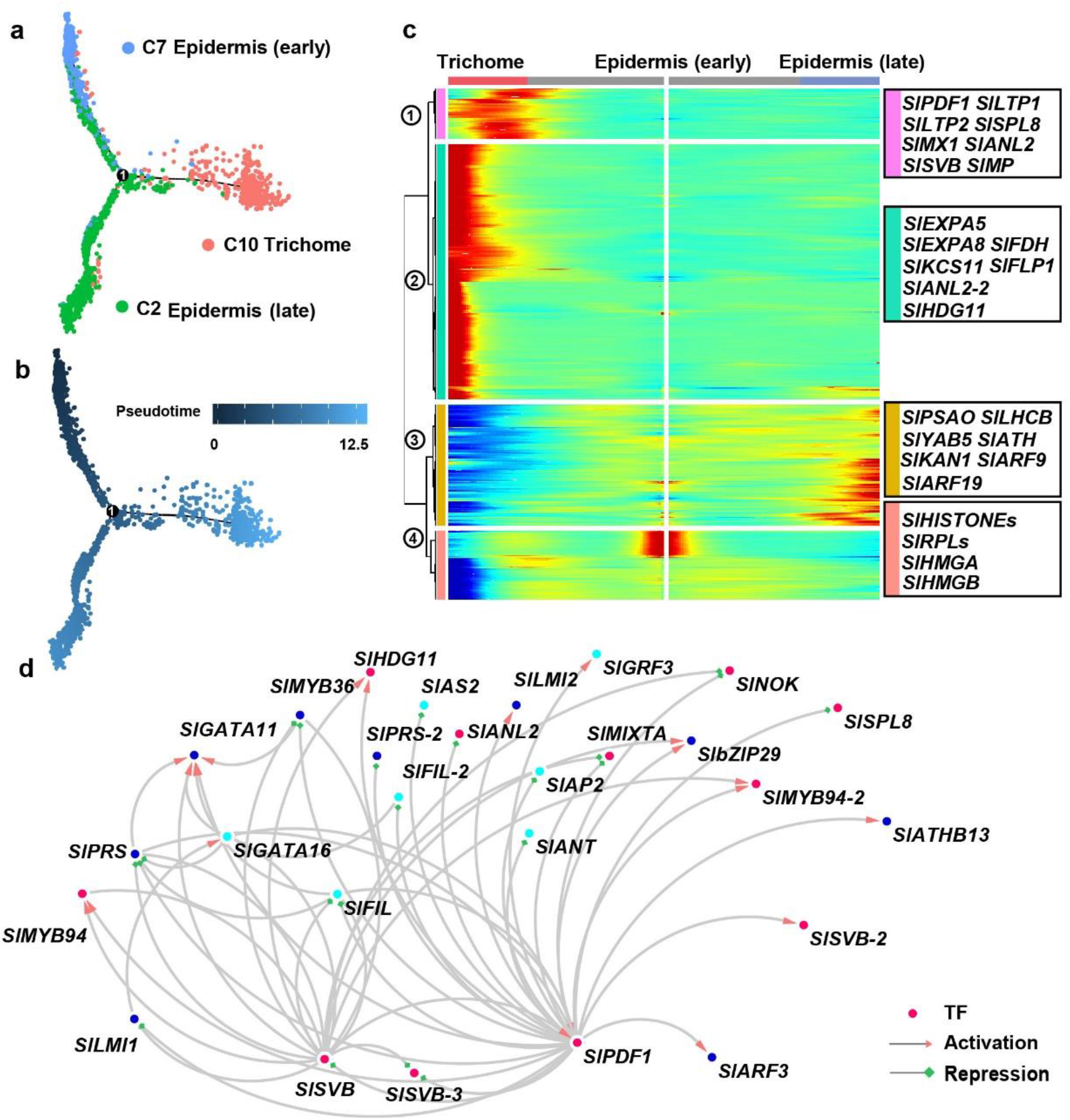
Development and differentiation of epidermal cells, including trichome cells. **a** and **b**, Developmental trajectory of epidermal and trichome cells showing clusters (**a**) and pseudotime (**b**). **c**, Gene expression patterns of epidermal cell differentiation along the pseudotime. **d**, GRN underlying epidermal cell differentiation. Dots represent transcription factors, edges indicate regulatory relationships, in which arrows for activation and squares for repression. The colors of dots represent enrichment in clusters as shown in Extended Data Fig. 9.

The existence of cell walls significantly encumbered the application of single cell transcriptomics in plant research. In this study, we develop a tissue processing pipeline to enable snRNA-seq profiling of virtually any plant cell type. Furthermore, snRNA-seq is expected to alleviate ectopic gene expression changes associated with protoplasting. We have applied snRNA-seq to obtain a high-resolution cellular expression atlas of the tomato shoot. With thicker cell walls and thicker tissues, tomato shoot apices are more resistant to protoplasting than *Arabidopsis* shoot apices. Nevertheless, we were able to obtain nuclei representing diverse major cell types. Our snRNA-seq analysis identifies most of the known cell types and portrays the remarkable heterogeneity at the cellular and molecular levels, including trichome subtypes and other rare cell types. Additionally, we infer developmental trajectories of key cell types and insights into SAM organization and function. In summary, we provided a robust single-nucleus transcriptomic profiling pipeline, which can be widely applied to other species and tissues, and a valuable resource for the study of stem cell homeostasis and early organogenesis.

## METHODS

### Plant materials and growth condition

The tomato (*Solanum lycopersicum*) cv. M82 was used. The seeds were sterilized with 40% bleach and geminated on 1/2 MS medium with 1.5% phytagel in culture vessels at 23°C in long-day conditions (16 h light/8 h dark).

### Sample processing and nuclei preparation

Seedlings 2 weeks after gemination were dissected under a stereoscope. Shoot apices (SAM together with the first three primordia) were harvested and frozen immediately in liquid nitrogen and stored at −80°C until use. Because leaf development is continuous, samples with early P4 were occasionally included. Shoot apices were resuspended in 10 ml nucleus isolated buffer (NIB: 10 mM MES-KOH (pH = 5.4), 10 mM NaCl, 10 mM KCl, 2.5 mM EDTA, 250 mM sucrose, 0.1 mM spermine, 0.5 mM spermidine, 1 mM DTT) with protease inhibitor cocktail (0.1%) and homogenized using a homogenizer at a low speed on ice. After lysis on ice for 30 min, the homogenate was filtered throughout a three-layer nylon mesh (Calbiochem) twice. To eliminate chloroplasts, 10% Triton X-100 was added dropwise to the solution to a final concentration of 0.1% (v/v) until most chloroplasts were degraded. Then the nucleus suspension was centrifugated at 1000 *g* for 5 min. The pelleted nuclei were washed twice and then suspended in NIB buffer (Extended Data Fig. 3). The procedure of snRNA-seq was shown in Fig. 1a. The variability, integrity, and concentration of the nuclei were determined by trypan blue staining and counted under a microscope. The nuclei concentration was adjusted to ~1000 cells/μl with NIB and subject for encapsulation with the 10x Genomics Single cell cassette according to the manufacture’s instruction.

### snRNA-seq library construction and sequencing

Approximately 20,000 nuclei were loaded for encapsulation. The library was constructed according to the manufacture’s instruction using Chromium Single Cell 3’ Library and Gel Bead Kit v3. Sequencing was performed on the Illumina Novaseq6000 platform with 150 paired-end reads.

### Preprocessing of raw snRNA-seq data

A pool of 17,097 nuclei were obtained after prefiltration by Cell Ranger v3.1.0 (https://support.10xgenomics.com/single-cell-gene-expression/software/pipelines/latest/what-is-cell-ranger). ITAG4.0 reference genome and annotation files were downloaded from International Tomato Genome Sequencing Project (ftp://ftp.solgenomics.net/tomato_genome/annotation/ITAG4.0_release/). The mapping rate was 92.6% and sequencing saturation was 85.8%, indicating a high quality of the library.

### Bulk RNA-seq

Total RNA was extracted from shoot apices using an RNA extraction kit (Axygen). Library was constructed as described before^14,15^, and sequenced by Illuminated HiSeq in the 150-nt paired-end mode. Three independent biological replications were performed. After quality control, clean reads were mapped to tomato reference genome ITAG4.0 (ftp://ftp.solgenomics.net/tomato_genome/annotation/ITAG4.0_release/) with Tophat2^25^. The counts were extracted using HTSeq and RPKM was calculated by edgeR^26^. The correlation between the RPKM from the bulk RNA-seq and the snRNA-seq was visualized using ggplot2, and the Spearman correlation was calculated.

### Cell clustering and annotation

Before cell clustering, we further removed low-quality nuclei with detected genes less than 500 or more than 2000 by Seurat3 (v3.1.2)^27^. Nuclei with mitochondrial genes contributing to over 1% and chloroplast genes contributing to over 5% were also filtered out. The feature matrix obtained after the above filtration was sent for further analysis. For cell clustering, the data was normalized by “LogNormalize” and highly variable genes were calculated with “FindVariableFeatures” with the “vst” method. The data was reduced to ~50 PCs and evaluated by JackStraw and Elbow, which showed that 20 PCs contributed to the majority of differentiation. Then “RunPCA” was performed to do the linear dimensional reduction, with the setting npcs = 30. Cells were clustered by “FindNeighbors” and “FindClusters” using the first 20 dims with resolution as 0.7. Cluster marker genes were identified using “FindAllMarkers” with parameters logfc.threshold = 0.5 and min.pct = 0.25, which means that log2 fold change of average expression is more than 0.5 and minimum cell percentage for marker genes is more than 25%.

### Homologous gene annotation

To utilize gene function and expression knowledge obtained in *Arabidopsis*, we identified tomato homologs of *Arabidopsis* genes. BLASTP was performed using tomato proteins as query against *Arabidopsis* proteins. The best hit with an *e*-value lower than 1e^-15^ was retrieved as the homologous gene. The correspondence of tomato and *Arabidopsis* genes is provided in Extended Data Table 6.

### Comparison with cell type-specific transcriptomic data

The gene expression data for *Arabidopsis* vegetative shoot apex and inflorescence SAM domains were retrieved^14–17^. Domain-specific genes were identified by defining genes with at least 2 times over the average expression of all domains. Enrichment analysis was performed as previously described^15^.

### Construction of developmental trajectory

We carried out pseudotime analysis with Monocle2 package (v 2.10.1)^28^ to order cells along the developmental process. In brief, cell expression matrices with specific clusters were retrieved as input. The dataset was rescaled with “estimateSizeFactors” and “estimateDispersions” functions. Then the variance was calculated by “dispersionTable” and variable genes were found by “FindVariable”. The data was reduced to two components with “DDRTree”. Cells were ordered along the pseudotime by “orderCells” and the developmental trajectory was visualized using “plot_cell_trajectory”. Pseudotime-dependent gene expression patterns were visualized with “plot_pseudotime_heatmap” function. To identify key genes for the cell fate transition, we applied BEAM algorithm to analyze the branch-dependent differentially expressed genes and used “plot_genes_branched_heatmap” function for visualization. Cluster-specific genes, pseudotime-dependent genes, and branch-dependent genes were sent to agriGO for GO enrichment analysis, respectively^29^.

### GRN analysis

To illustrate the gene regulatory relationships, we extracted the expression information of transcription factors with differential expression patterns along pseudotime trajectories. Their pseudotime values were normalized between 0 and 1. Then GRN was inferred using SCODE^30^ with the parameter z set to 4. We repeated the simulations 50 times to obtain reliable relationships. GRNs were visualized in Cytoscape^31^.

### Reporting summary

Further information on research design is available in the Nature Research Reporting Summary linked to this article.

## DATA AVAILABILITY

The raw snRNA-seq data are available from the NCBI SRA database with BioSample accession number SAMN16069893.

## ACKNOWLEDGEMENTS

We thank Chuanyou Li and Lei Deng for assistance with tomato growth. The work was supported by the Strategic Priority Research Program of CAS grant no. XDA24020203 to Y.J., and by a National Natural Science Foundation of China (NSFC) grant no. 31970805, 31961133010, and a Youth Innovation Promotion Association of CAS award no. 2017139 to C.T.

## AUTHOR CONTRIBUTIONS

C.T. and Y.J. conceived the project and designed experiments. C.T. and Q.D. performed experiments and analyzed data. M.X. and F.D. performed experiments. C.T. and Y.J. wrote the manuscript with inputs from all authors.

---

**Extended Data Fig. 1:**
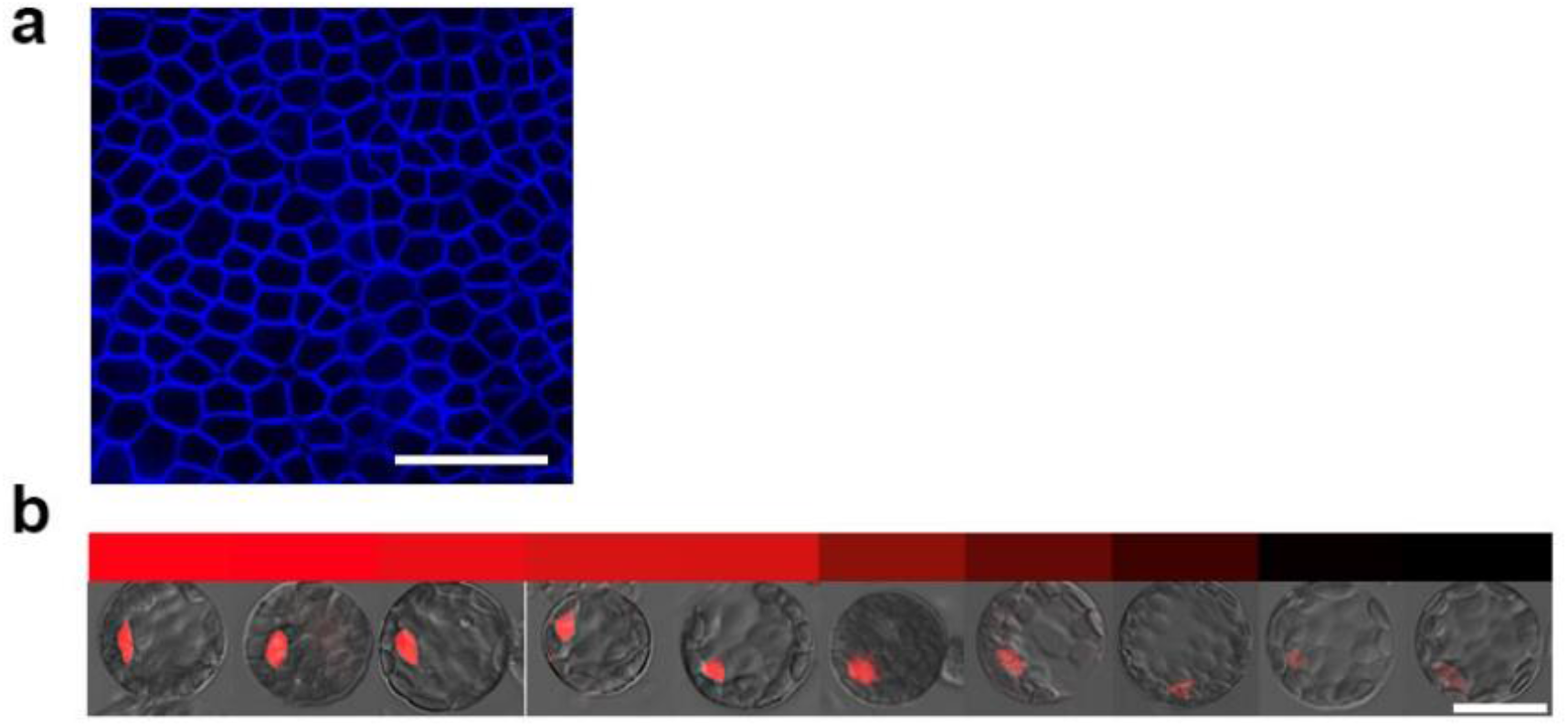
Gene expression pattern of *WOX2* gene in leaf and protoplasts. Confocal images of *Arabidopsis pWOX2::NLS-DsRed* plants with DsRed signals red and cell walls stained with FB28 (blue). a, There are no DsRed signals detected in all leaf cells, including epidermis and mesophyll cells. Bar = 50 μm. b, DsRed signal is frequently (21.3%) detected in leaf-derived protoplasts with variable expression levels. Representative protoplasts are shown. The color bar above shows fluorescence intensity quantification. Bar = 25 μm.

**Extended Data Fig. 2:**
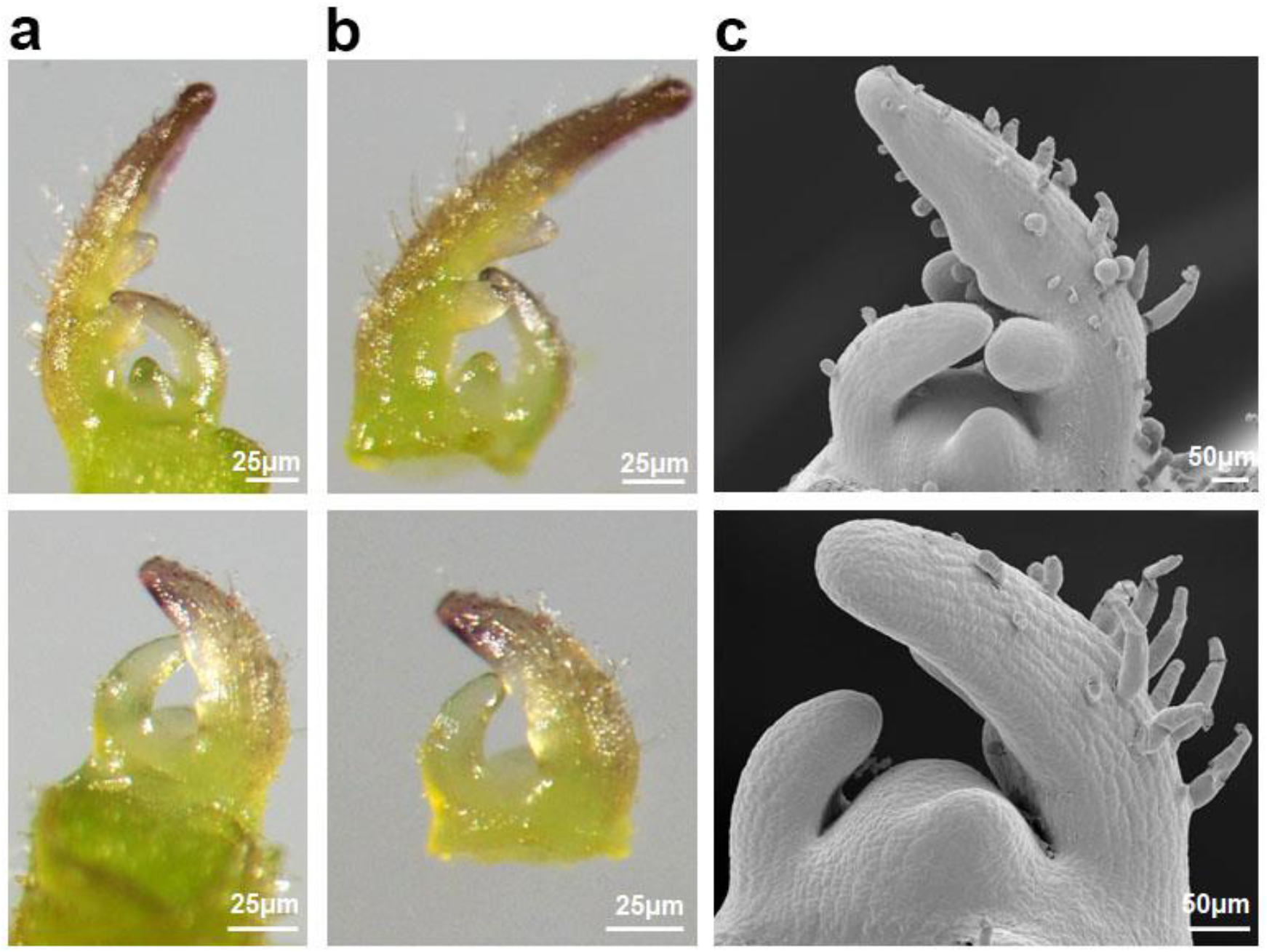
The anatomical structure of tomato shoot apex used for snRNA-seq. **a**, Shoot apices containing the SAM and early leaf primordia up to P_3_. **b**, Dissected shoot apices subject to nuclei isolation. **c**, Scanning electron microscopy photos of tomato shoot apices showing detailed cell morphology. The upper panel displays shoot apices with late P_3_ and the lower panel shows shoot apices with early P_3_.

**Extended Data Fig. 3:**
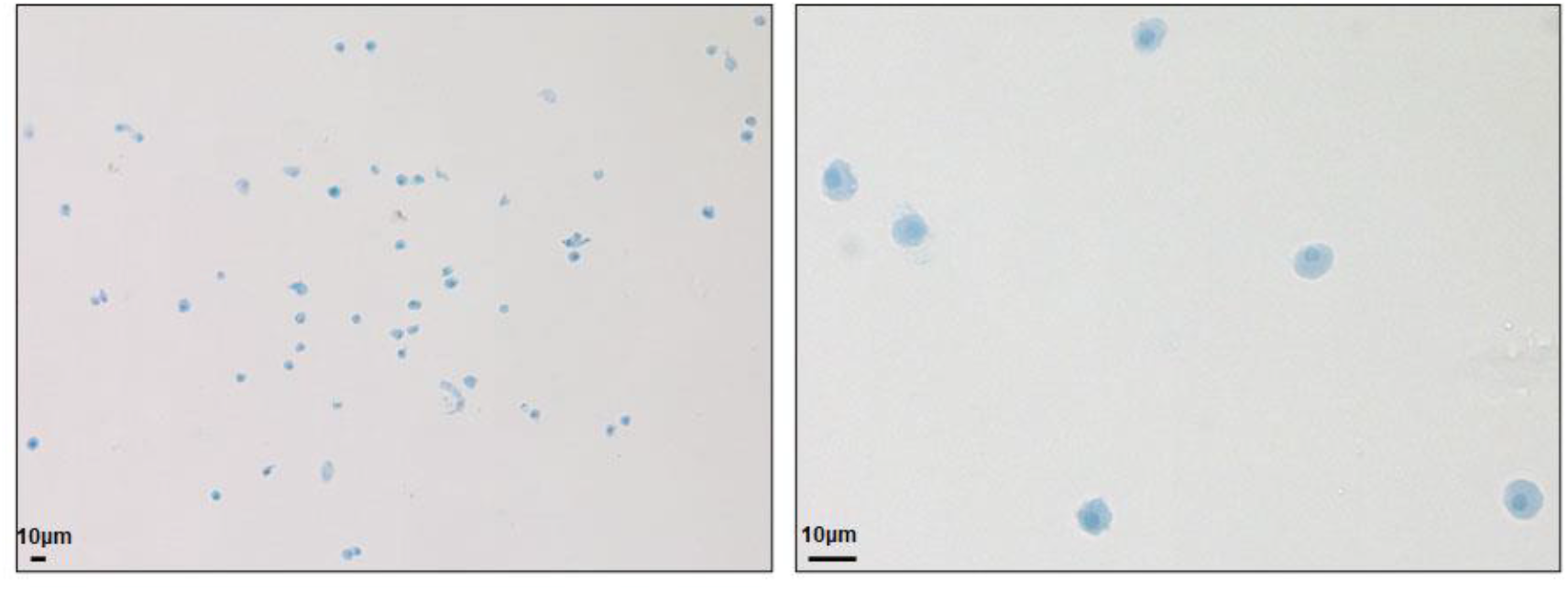
Isolated nuclei of tomato shoot apex stained with trypan blue.

**Extended Data Fig. 4:**
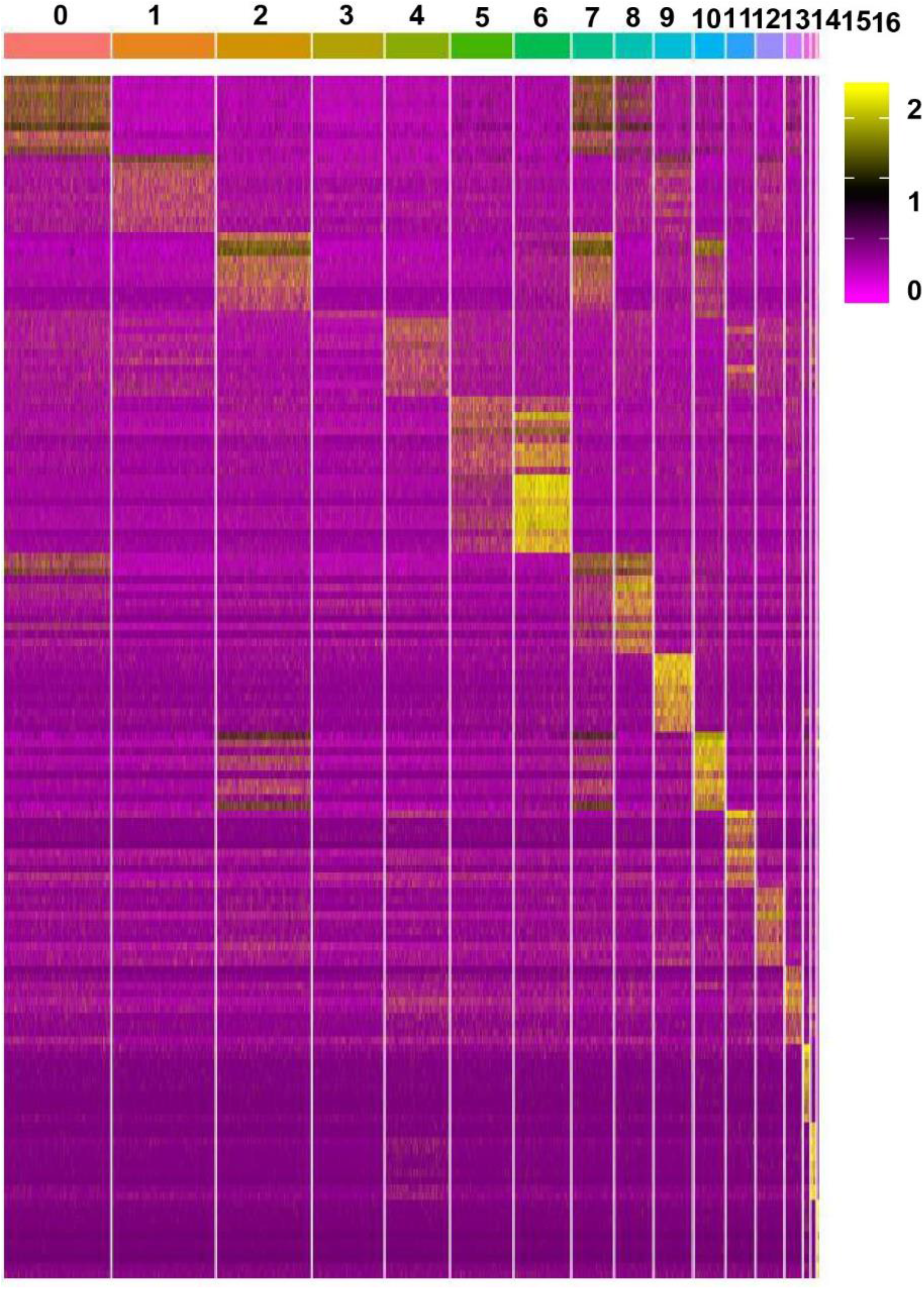
Heatmap of the expression of top 10 enriched genes of each cell cluster.

**Extended Data Fig. 5:**
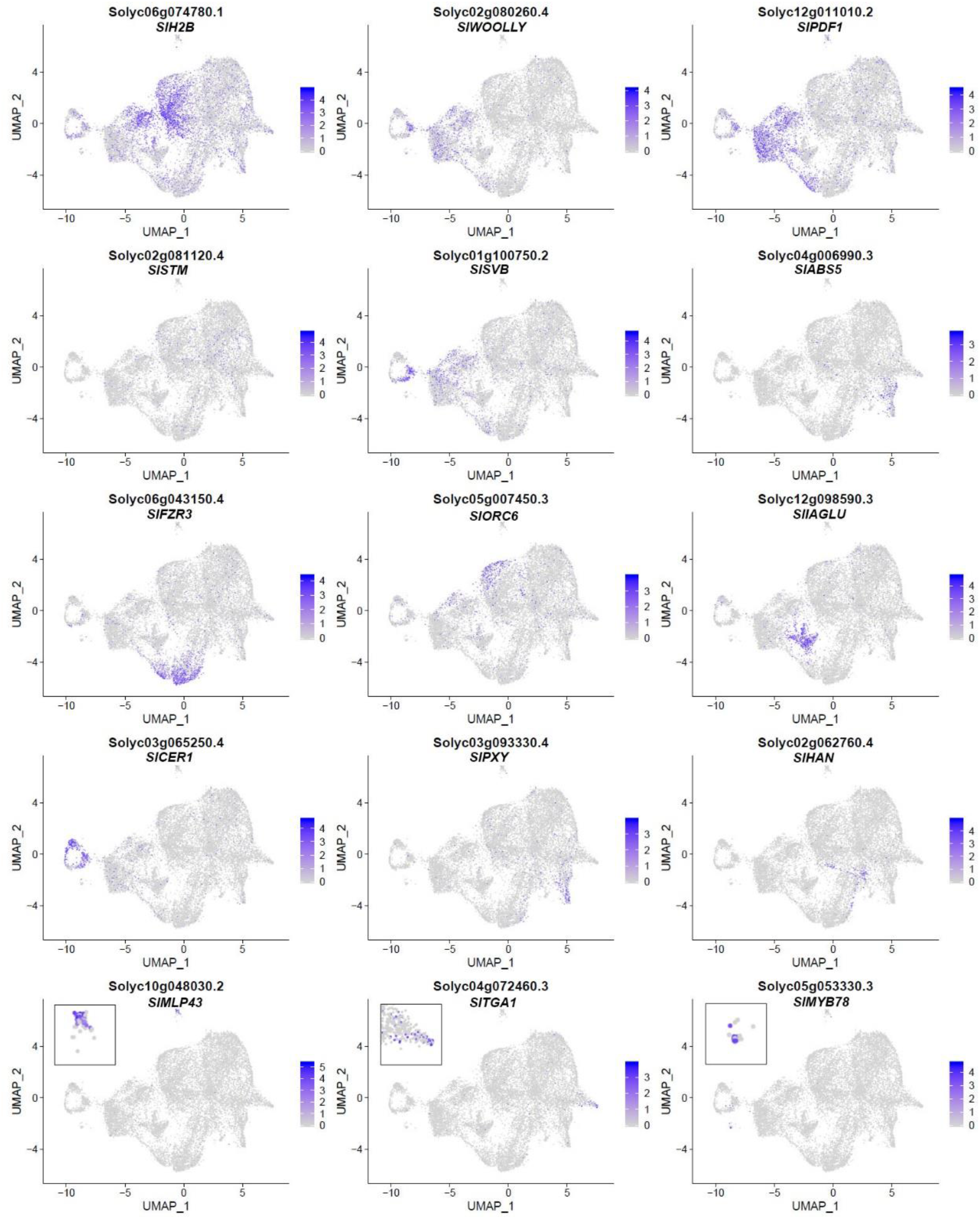
The distribution features of representative marker genes on uMAP clusters as shown in Fig. 2c.

**Extended Data Fig. 6:**
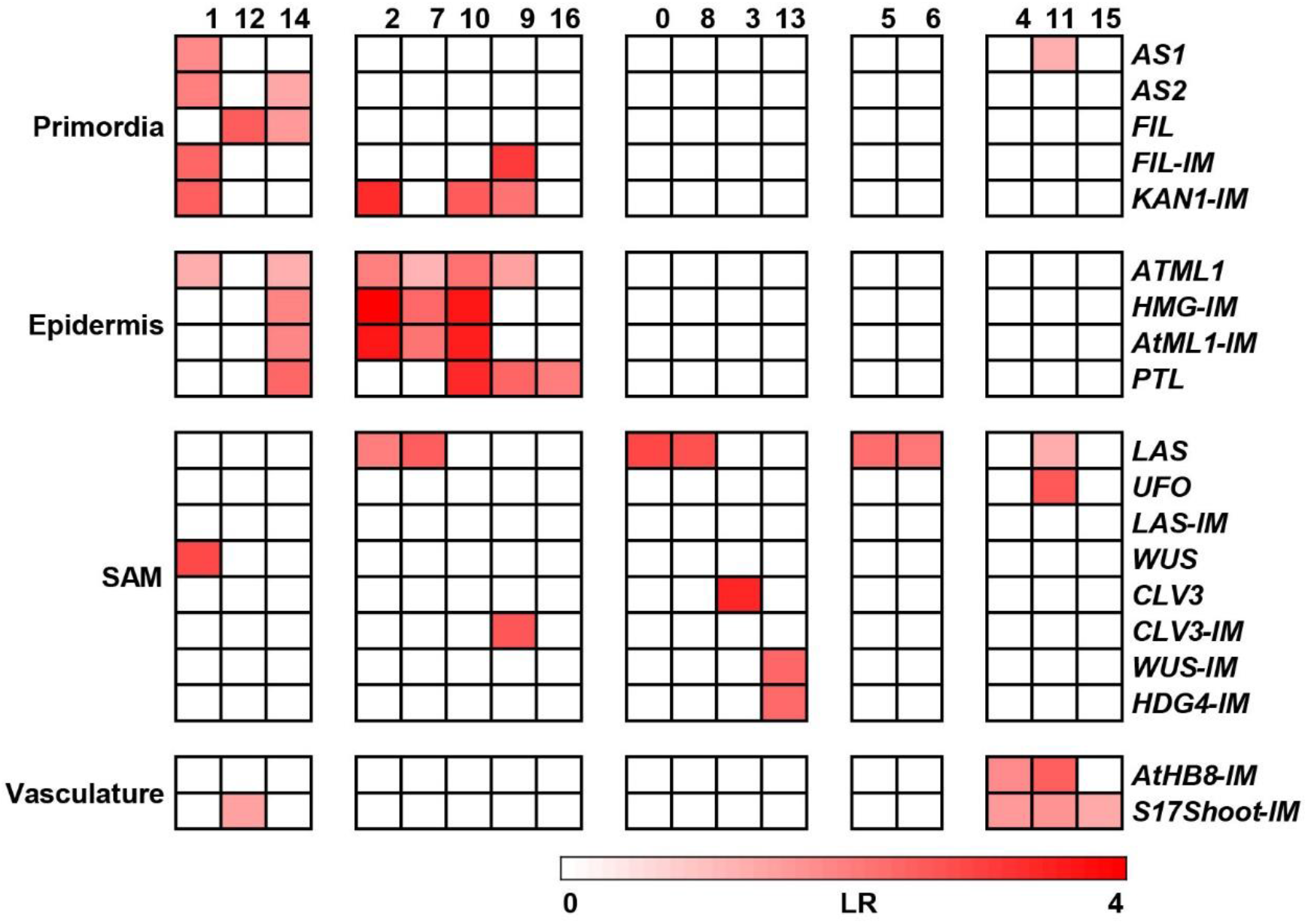
Enrichment analysis between cluster-enriched genes and cell type-specific genes. Published *Arabidopsis* vegetative shoot apex and inflorescence meristem cell typespecific genes are used, including genes enriched in the following cell types: AS1 (young primordia, 2342 genes), AS2 (leaf adaxial, 1323 genes), ATML1 (epidermis, 1989 genes), CLV3 (central zone, 2119 genes), FIL (leaf abaxial, 856 genes), LAS (boundary, 1515 genes), PTL (leaf margin, 580 genes), UFO (peripheral zone, 311 genes), WUS (organization center, 476 genes), AtHB8-IM (shoot xylem, 2079 genes), CLV3-IM (Central zone, inflorescence, 314 genes), FIL-IM (organ primordia, inflorescence, 382 genes), HMG-IM(Meristematic L1 layer, 595 genes), AtML1-IM (epidermis, inflorescence, 977 genes), HDG4-IM (subepidermis, inflorescence, 305 genes), WUS-IM (OC, inflorescence, 298 genes), LAS-IM (adaxial organ boundary, inflorescence, 372 genes), KAN1-IM (Abaxial organ boundary, inflorescence, 563 genes), and S17Shoot-IM (shoot phloem, 2421 genes). Cell clusters identified by snRNA-seq are shown as columns, and cell type-specific genes are shown as rows. Levels of enrichment were quantified by log2 odds ratio (LR), and colored accordingly.

**Extended Data Fig. 7:**
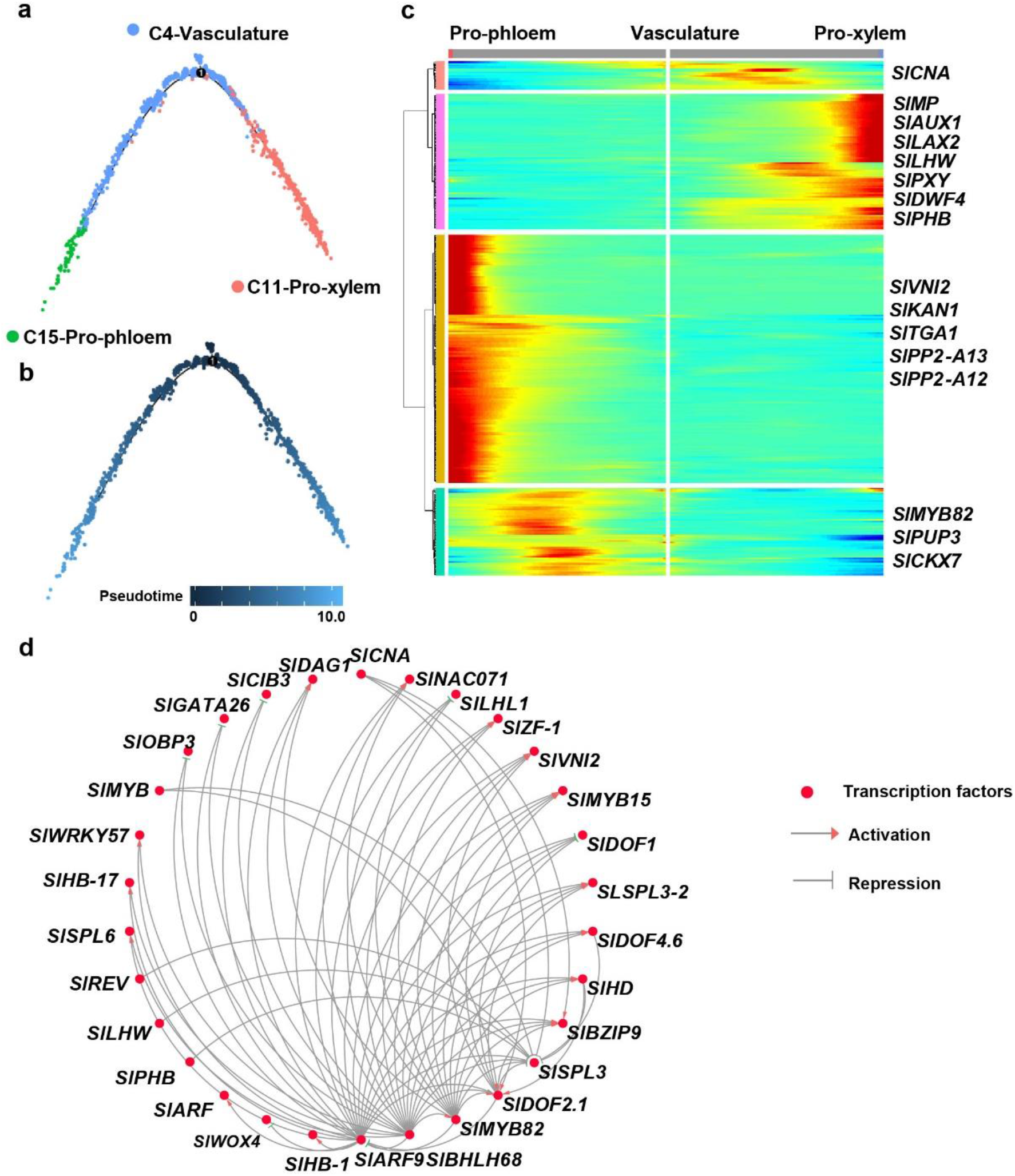
Developmental trajectory of vasculature cells. **a** and **b**, Developmental trajectory of vasculature cell differentiation showing clusters (**a**), and the pseudotime (**b**). **c**, A heatmap showing branch-dependent gene expression patterns. Putative key regulatory genes are shown in the right panel. **d**, GNR involved in xylem and phloem differentiation.

**Extended Data Fig. 8:**
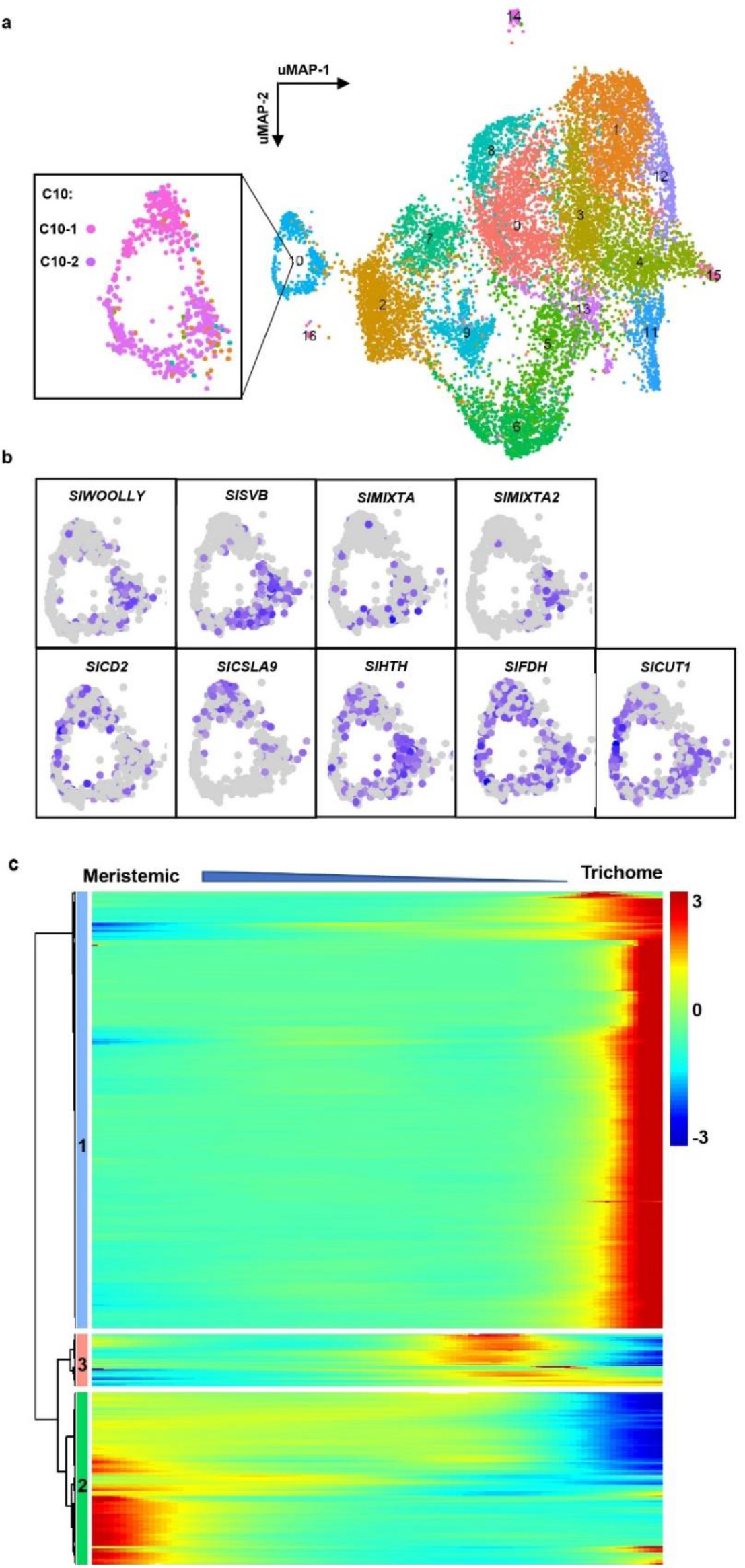
Subclusters of trichome cells and differential gene expression patterns over pseudotime. **a,** Subclusters of trichome cells on uMAP clusters as shown in Fig. 2c. **b,** Differential gene expression in trichome subclusters. **c,** A heatmap displaying differential gene expression patterns over pseudotime along trichome differentiation.

**Extended Data Table 1. Cluster-specific genes.**

**Extended Data Table 2. Correspondence between gene ID and gene names.**

**Extended Data Table 3. GO enrichment of cluster-specific genes.**

**Extended Data Table 4. Go enrichment of trichome subcluster genes.**

**Extended Data Table 5. GO enrichment of branch-dependent genes over trichome differentiation.**

**Extended Data Table 6. Correspondence between tomato and *Arabidopsis* homologous genes.**

## Notes

### Competing Interest Statement

The authors have declared no competing interest.

